# Gradient boosting regression and convolution improve deconvolution of bulk transcriptomes

**DOI:** 10.64898/2026.03.03.709368

**Authors:** Maik Wolfram-Schauerte, Thomas Vogel, Laura Achauer, Sara Maria Fälth Savitski, Hanati Tuoken, Eric Simon, Kay Nieselt

**Author notes:** Corresponding authors. Maik Wolfram-Schauerte, Eric Simon, Kay Nieselt.

## Abstract

Bulk cell type deconvolution aims to estimate cell type composition from bulk transcriptomic data. So-called pseudobulk simulation, where single-cell RNA-seq data is aggregated to bulk-like expression profiles, represents a central concept in training and testing of deconvolution tools. However, deconvolution methods often lack interpretability and struggle to generalize from simulated to real data. We present GrooD (GradientBoostedDeconvolution), a second-generation deconvolution tool that uses gradient boosted trees trained on pseudobulks from scRNA-seq references. GrooD’s pseudobulk simulations account for donor and condition variability to better model the complexity of real-world transcriptomes. We show that GrooD achieves state-of-the-art and superior deconvolution performance on human blood biospecimen. Furthermore, Grood provides visualizations of feature loadings and deconvolution results, that allow mechanistic insights for biological interpretation. We further integrate a convolution framework to assess the transcriptomic similarity between bulk and pseudobulk data, showing that higher similarity can indicate better deconvolution performance. GrooD deconvolution of blood transcriptomes from a large sepsis patient cohort identifies meaningful shifts in immune cell type composition that are associated with disease severity. By combining interpretability, robustness, and heterogeneous pseudobulk simulation, GrooD represents a powerful, user-friendly second-generation tool for cell type deconvolution.

## Introduction

Cell type-specific gene expression and cell type abundance, state and dynamics are key factors in health and disease. Single-cell RNA sequencing (scRNA-seq) has made this information increasingly accessible, offering unprecedented insight into cell type heterogeneity. However, scRNA-seq remains low-throughput, cost-intensive, and does not reliably quantify cell type proportions in complex tissues. Methods like fluorescence-activated cell sorting (FACS) and immunohistochemistry (IHC) allow direct quantification of cell types but similarly lack scalability. In contrast, bulk RNA sequencing (bulk RNA-seq) is a cost-effective, high-throughput technique widely used in large clinical studies. Yet, bulk RNA-seq masks underlying cell type proportions and gene expression, limiting its resolution.

To overcome this, computational deconvolution methods have been developed to infer cell type composition and expression from bulk RNA-seq, reviewed in [14]. These methods can be distinguished into first- and second-generation methods. Second-generation approaches incorporate scRNA-seq references and leverage a variety of machine learning strategies, including regression [7, 18, 11, 13] (e.g., MuSiC [19]), probabilistic (e.g., BayesPrism [4]), and neural network models (e.g., Scaden [12], TAPE [3]). Despite progress, several challenges remain [20, 5, 14, 1, 17]: 1) Evaluation often relies on pseudobulks - synthetic bulk-like profiles generated by aggregation of single-cell expression profiles from scRNA-seq data with associated cell type proportions based on the number of sampled cells per cell type. Since generated from scRNA-seq data, pseudobulks are more accurately deconvoluted than real bulks using second-generation methods [6]. 2) Interpretability remains limited, particularly for © probabilistic and neural network-based models [4, 12, 3]. 3) Also, most tools ignore unknown content - cell types absent from the reference, accounted for only by few methods [7]. 4) Recent work has also highlighted the lack of a holistic framework that integrates both deconvolution and its inverse, convolution [20]. While pseudobulk simulation is used to approximate convolution [6, 9], these simulations are used to benchmark deconvolution but rarely validated against real bulk data [20].

In this study, we introduce GradientBoostedDeconvolution (**GrooD**), a second-generation deconvolution tool addressing both interpretability and the lack of a holistic approach combining convolution with deconvolution. GrooD uses gradient boosted regression trained on heterogeneous pseudobulks generated from scRNA-seq data, simulating more realistic bulk profiles by accounting for cellular origin [9]. We further incorporate convolution into the analysis workflow, showing that the choice of the scRNA-seq reference significantly affects deconvolution. This improves deconvolution of real bulk RNA-seq data of peripheral blood mononuclear cells (PBMCs). We have applied GrooD to a sepsis cohort of over 300 individuals, recovering biologically meaningful cell type changes associated to patient subgroups specified by clinical outcome and explain these changes on gene expression level using its interpretability. Overall, GrooD includes a convolution perspective for deconvolution, thereby enhancing both performance and interpretability.

## Materials and Methods

### Concept and implementation of GrooD

Overall, GrooD operates in four phases: pseudobulk simulation, model training, inference, and evaluation. In the first phase, GrooD receives the input data, namely a single-cell reference and bulk RNA-seq to be deconvoluted with optional ground-truth cell type proportions. Pseudobulks are then simulated from the single-cell reference. Pseudobulks can be generated with or without preserving the origin of individual cells (e.g., patient or condition), a feature we refer to as heterogeneous pseudobulk simulation. In the second phase, GrooD is trained on these pseudobulk expression profiles. Specifically, GrooD uses a gradient boosted tree model per cell type to predict its proportion from the entire pseudobulk expression vector. This allows GrooD to identify the most informative genes for each cell type in a bulk-like context. In the third phase, the trained model infers cell type proportions from independent bulk RNA-seq data. Finally, predictions are visualized and optionally evaluated against ground-truth proportions. The following sections detail these steps.

### GrooD modes and input data

Input for GrooD is a bulk RNA-seq dataset, normalized to transcripts per million (TPM), for which a deconvolution is sought, and optionally ground-truth cell type proportions. For training, GrooD accepts a single-cell reference dataset, either as raw single-cell count matrices or precomputed pseudobulks with corresponding cell type proportions. In *train test* mode, GrooD simulates pseudobulks, normalizes them to counts per million (CPM) or another selected normalization (*log*_2_(CPM) or rank), trains the model, and evaluates its performance. In *inference* mode, GrooD deconvolutes input bulk RNA-seq data and optionally evaluates predictions if reference proportions are available. The *all* mode combines pseudobulk simulation, training, inference, and evaluation in a single workflow. When using this mode, the intersection of genes annotated in bulk and single-cell data can be used to avoid training on genes annotated in the reference but not in the bulk. We recommend consistent gene reference annotation for single and bulk RNA-seq data.

### Heterogeneous pseudobulk simulation

GrooD’s pseudobulk simulation extends the simulator implemented in Scaden [12, 3] by allowing for multi-threading to accelerate computation, heterogeneous pseudobulk simulation and various count normalization options. By default, pseudobulks are generated using random cell type proportions that sum to one, with random cell types randomly omitted (set to zero) per pseudbobulk. Alternatively, user-defined proportions can be supplied for simulation. Based on these proportions, single cells are sampled from the cell type-annotated single-cell reference per cell type (AnnData object must contain `cell type` column in observations). Counts of single-cells are aggregated and scaled to counts per million (CPM). Optional *log*_2_- or rank-based normalization can be applied. Each pseudobulk is paired with its corresponding cell type proportions. By default, GrooD simulates 1000 pseudobulks using 1000 cells each.

To enable heterogeneous simulation, single-cell-level metadata can be used to restrict sampling. The `individual` field indicates the donor/individual a single-cell was derived from, while `condition` can denote experimental batch or phenotype background of the patient a cell was derived from (e.g., healthy vs. diseased). During heterogeneous simulation, only cells from a specific condition or individual are selected per pseudobulk.

### GrooD training

GrooD uses gradient boosted trees for deconvolution, for which the `GradientBoostingRegressor` from scikit-learn, wrapped in a `MultiOutputRegressor` to train separate models for each cell type, is employed. By default, each regressor uses 500 trees (maximum depth 4), a minimum of 5 samples per split, and a learning rate of 0.01, optimized with an L2 loss. Training is performed on 80% of the pseudobulks, while the remaining 20% are held out for validation. Evaluation includes regression plots, error metrics, and cell type-specific feature importance based on mean decrease in impurity (MDI). The top 20 genes per model are extracted to assess key contributors to predictions.

### GrooD inference and evaluation

During inference, TPM-normalized bulk RNA-seq data are subset to the gene set used in training and normalized accordingly to the train data. GrooD predicts cell type proportions per bulk sample. Importantly, inferred cell type proportions are scaled to a total sum of 1 only, if their sum exceeds 1. Thereby, predicted proportions remain in a biologically meaningful range, while GrooD allows to predict less than 100% proportions leaving a potential residual for unknown content. In addition, GrooD visualizes predicted cell type proportions and optionally evaluates against the ground-truth proportions using correlation and error metrics (Supplementary Information).

### Data and deconvolution assessment

For GrooD assessment and benchmarking, we focussed our analysis on blood as the biospecimen of interest, for which most suitable datasets are available [20, 5].

### Single-cell references and pseudobulk generation

Single-cell data for *Hao* were retrieved from GEO (GSE164378, PBMC CITE-seq RNA 3P) [8]. The *Steixner-Kumar* dataset (GSE267033) was downloaded and processed similarly [16]. Cells with *<* 500 detected genes and genes expressed in *<* 40 (Hao) or *<* 30 (Steixner-Kumar) cells were excluded. Cell type annotations in *Hao* were aligned with *Finotello*, while *Steixner-Kumar* annotations were grouped into monocytes, lymphocytes, and neutrophils to match the *OConnell* dataset. All preprocessing was performed with scanpy and annotations were visually validated. Single-cell data were split 80/20 into stratified train/test sets.

Using the *Hao* test set, 50 pseudobulks were generated with random cell type proportions to evaluate different simulation strategies (canonical, batch-specific, patient-specific). Each pseudobulk contained 500 sampled cells, aggregated and normalized to CPM. To compare pseudobulks with real bulk data, an additional set of pseudobulks was simulated from the *Hao* train set, using either the cell type proportions given by the single-cell data or those derived from matching bulk samples. CPM-normalized pseudobulks were then compared to TPM-normalized bulk samples through correlation (Supplementary Information).

### Bulk RNA-seq data and deconvolution

GrooD was evaluated using default settings (see training and pseudobulk simulation). Pseudobulks were simulated from raw single-cell counts, CPM-normalized, and used to train GrooD with default model parameters. Runs were performed in *all* mode using protein-coding genes. Input bulk data were TPM-normalized.

TPM-normalized bulk RNA-seq data from the *Finotello* dataset were retrieved from GEO (GSE107572) [7] and associated ground-truth cell type proportions were downloaded (https://figshare.com/articles/dataset/Validation_real/25347757) [5]. Neutrophils and “Other” cell types were excluded. Deconvolution of the *Finotello* data was performed using the *Hao* train set. Raw bulk RNA-seq data from the *OConnell* dataset (SRP429744) [15] were processed using a Nextflow pipeline including adapter trimming (cutadapt), mapping (HISAT2 to RefSeq GCF 000001405.40), and quantification (featureCounts). One sample with *<* 1*M* reads was discarded, and TPM-normalization was applied. Cell type proportions were extracted from SRA metadata, and eosinophils were excluded. *OConnell* deconvolution was informed by the *Steixner-Kumar* train set. The *Baghela* dataset (GSE185263) [2] was processed similarly. Deconvolution of the *Baghela* dataset was performed with GrooD trained on the *Hao* train set at default settings (see above).

### Benchmarking settings

Benchmarking against state-of-the-art tools Scaden [12, 3], BayesPrism [4] and MuSiC [19] was performed using a custom Nextflow pipeline. Bulk TPM-normalized data with matched ground-truth proportions and raw single-cell counts were used as input. All tools were run with default settings. Deconvolution performance was evaluated per sample and per cell type using correlation and error metrics (Supplementary Information).

## Results

Performance of GrooD on pseudobulk and bulk RNA-seq data As pseudobulk simulation remains the standard approach for benchmarking deconvolution tools, we first assessed GrooD’s performance on TPM-normalized pseudobulks simulated from a 20% subset of the *Hao* single-cell reference, comprising seven major PBMC cell types [8]. GrooD was trained on the remaining 80% using canonical pseudobulk simulation, without accounting for donor or condition origin. The model achieved high performance across all seven cell types (Fig. 1A), demonstrating that gradient boosted regression can accurately infer cell type proportions from transcriptomic data. This is in line with results from *Kassandra*, which also employs gradient boosted trees but is trained on real bulk data [21]. Unlike GrooD, however, *Kassandra* does not leverage pseudobulks and is limited to predefined cell types.

**Fig. 1:**
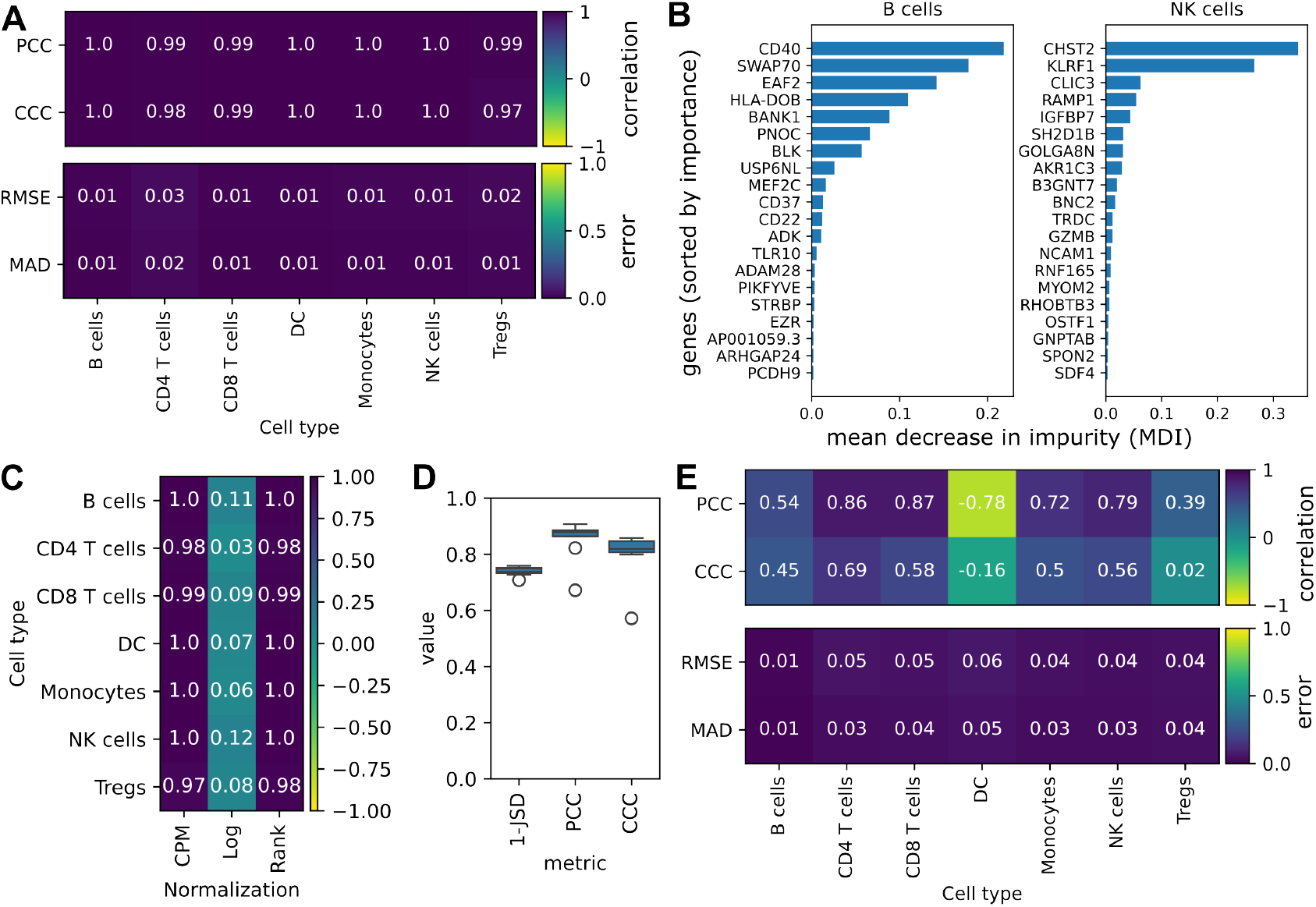
Basic performance of GrooD on gold-standard PBMC data. **A**) Per cell type deconvolution performance of GrooD on PBMC pseudobulk data (n=50) across multiple quality metrics. **B**) Feature importances (mean decrease in impurity (MDI)) for B cells and NK cells learned by GrooD, visualized as bar plots. **C**) Impact of input data normalization (CPM, log, rank) on GrooD’s deconvolution performance on PBMC pseudobulks (n=50), evaluated per cell type using concordance correSlation coefficient. **D**) Similarity between pseudobulks simulated from the *Hao* reference and bulk RNA-seq samples from the *Finotello* dataset in a convolution-like setting, wherein the same proportions from Finotello are used for pseudobulk simulation. Similarity is measured by CCC, 1-JSD and Pearson correlation on CPM-/TPM-normalized gene expression level per sample (n=9). Deconvolution performance of GrooD on the *Finotello* bulk dataset (n=9) using the *Hao* reference. CCC: concordance correlation coefficient, PCC: Pearson correlation coefficient, RMSE: root mean squared error, MAD: mean absolute deviation, JSD: Jenson-Shannon Divergence. Cell types: NK cells: natural killer cells, DC: dendritic cells, Tregs: T regulatory cells.

To evaluate data similarity between train and test settings, we implemented a convolution control by simulating pseudobulks with matched cell type proportions. Since all pseudobulks were derived from the same dataset, we expected - and observed - high similarity (Supplementary Fig. S1A).

One advantage of GrooD is its decision-tree-based approach. Therefore, we examined GrooD’s feature importance to assess biological interpretability. For each cell type, GrooD identified canonical marker genes, such as *CD8B* for CD8 T cells and *KLRF1* for NK cells, without explicit prior marker gene information (Fig. 1B). This highlights GrooD’s capacity to identify meaningful biological signals directly from the bulk expression context.

By default, GrooD simulates pseudobulks from raw single-cell counts and normalizes them to CPM, aligning them with the scale of TPM-normalized bulk data. To evaluate how normalization affects accuracy, we also tested log2-transformation and rank-normalization for both train and test data. CPM outperformed log-normalization and performed comparably with rank-normalization (Fig. 1C), indicating GrooD sensitivity to data normalization and CPM as a suitable normalization strategy.

Next, we applied GrooD to real bulk RNA-seq data from the *Finotello* dataset [7], which includes transcriptomes from nine individuals with matched cell type proportions measured by FACS. While the dataset contains eight cell types and a residual (“Other”), the *Hao* reference includes seven overlapping PBMC cell types, which together represent 90% of the measured cell types. Hence, we restricted the evaluation to these seven cell types.

To assess the suitability of the *Hao* reference for deconvolution, we simulated nine pseudobulks with matching cell type proportions and compared them to the *Finotello* bulk transcriptomes. This convolution-like comparison showed that restricting both datasets to common protein-coding genes improved pseudobulk–bulk similarity (Fig. 1D, Supplementary Fig. S1B,C). Therefore, we used protein-coding intersecting genes for GrooD deconvolution.

GrooD recovered meaningful cell type proportions in the *Finotello* bulk data (Fig. 1E). Predictions correlated well with FACS measurements for major cell types and maintained low error rates even for less abundant populations. However, certain low-abundance types (e.g., dendritic cells (DCs), T regulatory cells (Tregs)) showed reduced or even negative correlation, indicating over- or underestimation. Nonetheless, prediction errors remained low overall. At the sample level, GrooD’s predicted compositions closely resemble the ground-truth (Supplementary Fig. S1D), supporting its generalizability from pseudobulk to real bulk RNA-seq.

In summary, GrooD delivers accurate and interpretable deconvolution from scRNA-seq-derived pseudobulks and performs well on real-world bulk data, even with imperfect reference overlap.

## Heterogenous pseudobulk simulation increases GrooD’s deconvolution performance

So far, we used GrooD in settings, where the origin of single-cells is not considered for pseudobulk simulation. However, recent work has shown that heterogeneous pseudobulks, which account for individual-origin or condition (e.g. disease state), show better the limitations of deconvolution tools than canonical pseudobulks [9], which we used for training and evaluation as described above. Therefore, we introduced heterogeneous pseudobulk simulation into GrooD training, aiming to train GrooD in a more realistic, challenging bulk context. For this, we trained three GrooD models: one with canonical pseudobulks, one with pseudobulks accounting for individual origin and one with pseudobulks accounting for batch origin as condition. Testing GrooD on pseudobulks generated from held out single-cell data, we observed that heterogeneous pseudobulks are less accurately deconvoluted by GrooD trained with canonical pseudobulks (Fig. 2A). In contrast, condition-specific simulation seemed to yield consistently higher performance across test pseudobulks.

**Fig. 2:**
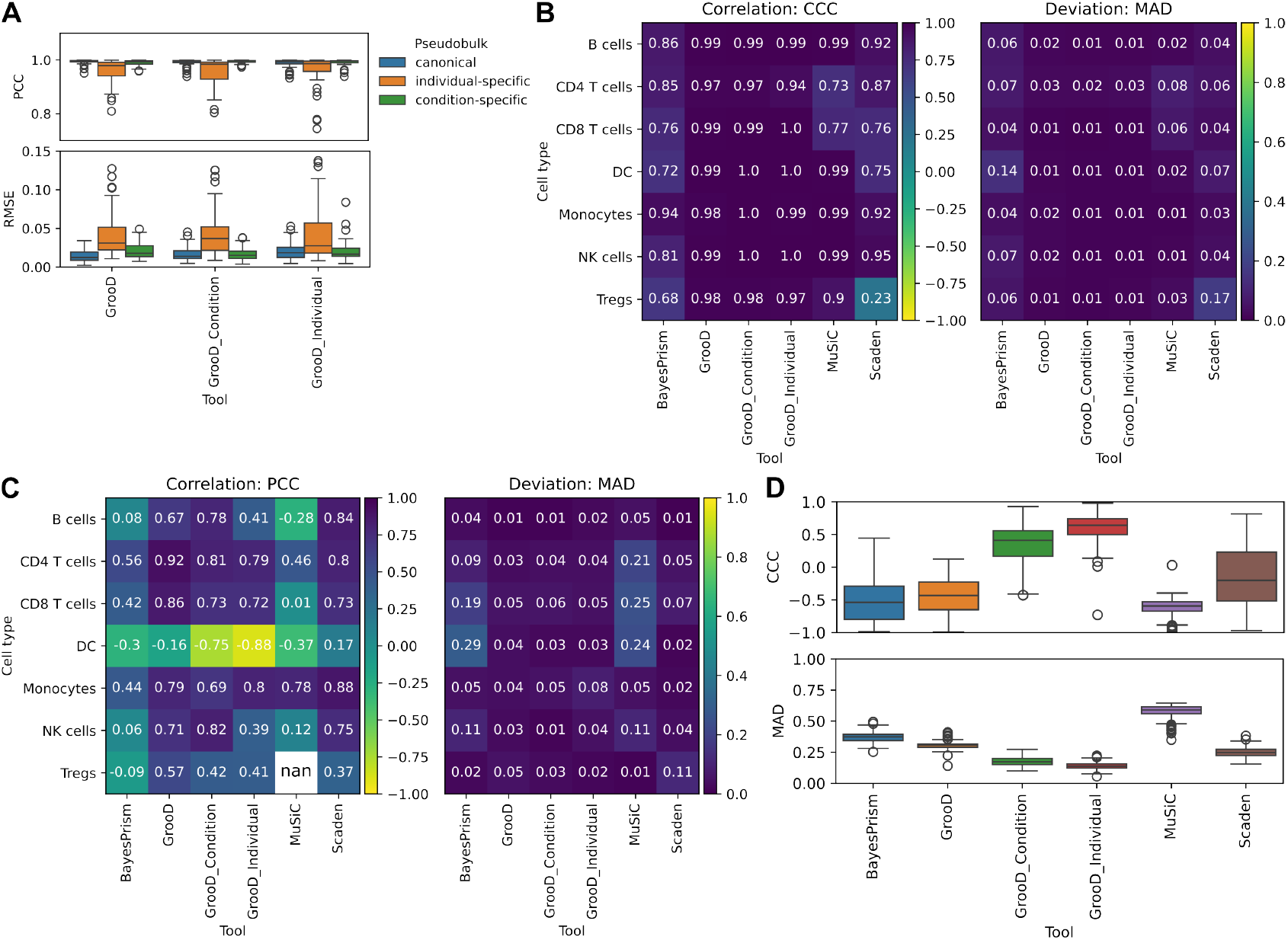
GrooD training with heterogeneous pseudobulks enhances bulk transcriptome deconvolution. **A**)Distribution of sample-wise correlation (PCC) and deviation (RMSE) from ground-truth cell type proportions for canonical, individual- and condition-specific pseudobulks (n=50 each) deconvoluted with GrooD. GrooD models were trained with either of the pseudobulk strategies. 80% of the *Hao* single-cell reference were used for training and pseudobulks were generated from the remaining 80% of cells.**B**) Cell type-wise deconvolution performance measured as CCC and MAD across three state-of-the-art deconvolution tools and GrooD in three different training modi w.r.t. pseudobulks. Performance was evaluated on n=50 heterogeneous pseudobulks simulated using random cell type proportions using 20% of single-cells from the *Hao* reference dataset. Residual 80% of the data used for training. **C**) Cell type-wise deconvolution performance on the *Finotello* dataset (n=9 bulk samples) measured by PCC and MAD using the *Hao* single-cell reference. No correlation measurement for Tregs for MuSiC, since MuSiC predicted fractions of 0.0 for all samples. **D**) Sample-wise deconvolution performance measured by CCC and MAD. Deconvolution was informed by the *Steixner-Kumar* single-cell reference and evaluated on three cell types (lymphocytes, monocytes, neutrophils) using the *OConnell* dataset (n=137 bulk samples). Error and correlation metrics as in Fig. 1.

To further evaluate GrooD’s deconvolution performance on heterogeneous pseudobulks, we compared GrooD against state-of-the-art tools, which have shown outstanding performance in recent benchmarks [5, 14], including BayesPrism [4], Scaden [12] and MuSiC [19]. We observed that GrooD outperforms the three tools for all cell types, both in correlation and error indicated by our benchmarking on condition-specific *Hao* pseudobulks (Fig. 2B). GrooD also showed superior performance, when trained with heterogeneous pseudobulks. We therefore speculated that this could enhance GrooD-mediated deconvolution of real bulk data.

We then used the *Hao* single-cell reference again to deconvolute the nine *Finotello* bulk samples and included GrooD trained with canonical and heterogeneous pseudobulks. We found that GrooD performed best across several cell types in both correlation and error metrics on the bulk data (Fig. 2B). Condition-specific pseudobulks improved especially prediction errors across several cell types, overall positioning GrooD at the top of the scale, even slightly outperforming Scaden in deconvolution of several PBMC cell types. It stood out that deconvolution of DCs was strongly anti-correlated for GrooD, in particular when trained in the individual-specific setting, while prediction errors were low.

Since the *Finotello* dataset comprises only nine samples, we sought to test a larger benchmarking dataset. We identified the *OConnell* dataset that measured the bulk transcriptomes of 137 human whole blood samples alongside the cell type proportions for four major cell types (monocytes, lymphocytes, neutrophils and eosinophils) [15]. Since we did not have a single-cell reference including eosinophils, we decided to exclude them from our evaluation. Using a suitable single-cell reference, *Steixner-Kumar* [16], we benchmarked GrooD against state-of-the-art tools on the *OConnell* dataset. Overall, all tools showed lower prediction accuracy on the *OConnell* dataset compared to the *Finotello* dataset (Fig. 2C). In order to explain this overall low performance, we compared the bulk and pseudobulks in a convolution-like setting, which revealed a very low concordance for all three metrics (Supplementary Fig. S2). Addressing this large variability by using a heterogeneous pseudobulk training, GrooD showed largely improved deconvolution performance (Fig. 2D).

## GrooD reveals cell type proportion trends in a sepsis cohort

Clinical studies often generate bulk RNA-seq data from large patient cohorts and provide associated phenotypic information, such as disease status or clinical outcome. Therefore, as a use case, we applied GrooD to a large sepsis cohort comprising 389 individuals [2], for which PBMC bulk transcriptomes and clinical outcome data are available. The cohort includes 44 healthy controls, 293 sepsis patients who survived, and 52 patients who died from sepsis. Using the *Hao* single-cell reference, we trained GrooD on canonical pseudobulks to predict the proportions of seven major PBMC types. We then predicted cell type proportions on the patient level from the bulk RNA-seq data and compared them in the three major patient segments (healthy vs. survived sepsis vs. died sepsis). Consistently, we observed clear, associated changes in particular cell types (Fig. 3A, Supplementary Fig. S3). Overall, we observed a reduction in the proportions of major immune cell types, including NK cells, CD4 T cells, and CD8 T cells, in sepsis patients, with increasing disease severity. Among these, CD4 T cells were most reduced in deceased sepsis patients (Fig. 3A). This agrees with the finding from the original paper that a subgroup of patients with an immune suppressive endo-phenotype shows high mortality and studies that highlighted decrease in T cell levels upon sepsis [10].

**Fig. 3:**
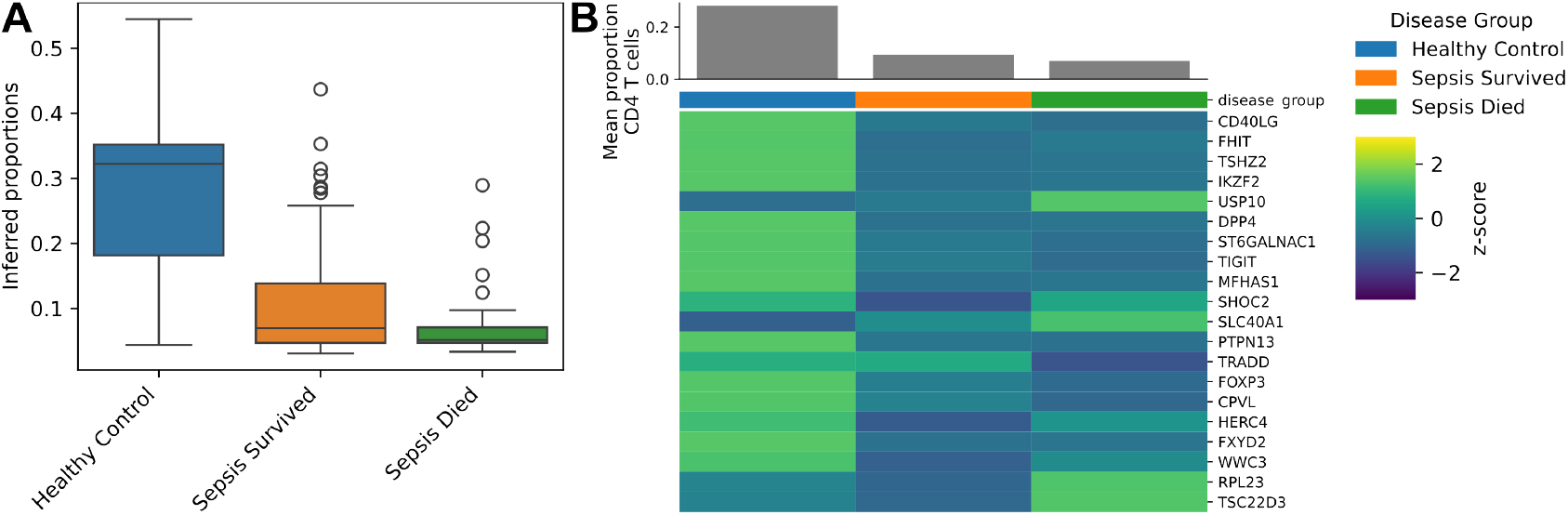
GrooD infers meaningful trends in cell type proportion changes in large patient cohorts. **A**) CD4 T cell proportions predicted with GrooD using the Hao reference [8] for 389 bulk samples from the Baghela dataset [2] per patient group (healthy control (n=44), sepsis survived (n=293), sepsis died (n=52)). **B**) Heatmap of z-score normalized mean expression per patient group of the 20 most important genes (sorted from most to least important from top to bottom) indicated by GrooD for predicting CD4 T cell proportions. The predicted mean proportions are indicated as bars on top of the heatmap.

GrooD is also very useful for biological interpretation because it provides the most relevant features (genes) per cell type and thus links bulk expression of individual genes to single cell types (Fig. 1B). We could use these marker genes as starting points for biomarkers to stratify specific patient segments as shown in Fig. 3B. Here, most marker genes were relatively low expressed in the subgroup of sepsis survivors, while healthy individuals generally show higher expression. Few marker genes tend to be expressed at elevated levels in deceased sepsis patients. Overall, this demonstrates that GrooD not only infers cell type proportions also from large cohort datasets, but can also aid in downstream analysis by identifying marker genes for each cell type, and thus allows interpretation of results and potentially patient segmentation.

## Discussion

In this study, we introduced GrooD, an interpretable, second-generation deconvolution tool based on gradient boosted trees. A key innovation of GrooD is its use of heterogeneous pseudobulk simulation during training, which significantly improves deconvolution performance on real bulk RNA-seq data compared to state-of-the-art tools. GrooD achieves competitive or superior results on both synthetic pseudobulks and real bulk datasets, suggesting that it is less biased toward pseudobulk-optimized settings. Importantly, GrooD maintains robust performance even when specific cell types are absent from the reference, demonstrating its potential to estimate unknown content - a common challenge in real-world bulk transcriptomics. We further demonstrate its utility in large patient cohorts by providing meaningful insights into shifts in immune cell proportions.

Our findings also underscore the need for holistic integration of single and bulk-based methods. We observed that deconvolution accuracy improves when convolution of the single-cell reference yields transcriptomic profiles closely resembling the bulk RNA-seq data. This supports previous observations that modeling convolution more accurately can enhance deconvolution performance [20]. GrooD implicitly addresses this by using heterogeneous pseudobulk simulation to better reflect biological and technical variability in real bulk data. Building on this, we plan to further develop convolution modeling strategies to support more generalizable and robust deconvolution approaches. While our benchmarking focused on blood-derived data, where high-quality reference datasets and ground-truth cell type proportions are available, GrooD has not yet been evaluated on solid tissues such as liver, lung, or brain. These tissues pose specific challenges due to increased cellular complexity, spatial heterogeneity, and limited availability of matched bulk and single-cell data. Future work will extend GrooD’s application to these tissues, although comparative benchmarking remains difficult due to the scarcity of suitable gold-standard datasets [20]. Here, single-nucleus RNA-seq might become a valuable additional reference type for GrooD.

In summary, GrooD advances current deconvolution approaches by combining interpretability, strong real-bulk performance, and a holistic integration of convolution principles. This represents a significant advancement of cell type deconvolution methods towards more accurate and decision guiding applications in clinical transcriptomics.

## Supplementary data

Supplementary Methods and Figures S1 - S3 are available in Supplementary Information.

## Data availability

GrooD is available at https://github.com/MaikTungsten/GrooD including code for all figures, analyses and processing steps relevant to this work. The pseudobulk simulator is also available as a standalone CLI tool https://github.com/MaikTungsten/PseudobulkSimulators. The benchmarking pipeline is available at https://github.com/MaikTungsten/Deconvolution_benchmarking. The RNA-seq pipeline is available at https://github.com/MaikTungsten/RNAseq_pipeline.

