## Supplementary Information for "Gradient boosting regression and convolution improve deconvolution of bulk transcriptomes"

Maik Wolfram-Schauerte, Thomas Vogel, Laura Achauer, Sara Maria Fälth-Savitski, Hanati Tuoken, Eric Simon and Kay Nieselt

February 2026

#### 1 Metrics

We used the **root mean squared error** (RMSE) for sample-wise (comparing cell type proportions or gene expression) and cell type-wise (only cell type proportions) comparisons. The RMSE is defined as

$$\text{RMSE} = \sqrt{\frac{1}{n}\sum_{i=1}^{n}(y_i - \hat{y}_i)^2}\ ,$$

where $y_i$ is the ground-truth value, i.e. the expression value of gene $i$ or proportion of a cell type $i$ in a sample, or the proportion for a cell type in a sample $i$. The number or genes or cell types is given by $n$. In addition, $\hat{y}_i$ represents the gene expression value of gene $i$ in a corresponding pseudobulk sample or the associated predicted cell type proportion of a cell type $i$ or sample $i$, respectively.

Similarly, we applied **mean absolute deviation** (MAD) both for sample-wise (comparing cell type proportions or gene expression) and cell type-wise (only cell type proportions) comparisons. The MAD is defined as

$$\text{MAD} = \frac{1}{n}\sum_{i=1}^{n}|y_i - \hat{y}_i|\ ,$$

where the notation is identical to RMSE.

We used two different correlation metrics, the Pearson and the concordance correlation coefficient, for the comparison of either bulk and pseudobulk expression profiles or predicted and ground-truth cell type proportions. The **Pearson**

**correlation coefficient** (PCC) describes the linear correlation of two paired samples. It is defined as

$$\text{PCC} = \frac{\sigma_{ij}}{\sigma_i \sigma_j}$$

with the covariance $\sigma_{ij}$ defined as

$$\sigma_{ij} = \sqrt{\frac{1}{n-1}\sum_{k=1}^{n}(x_k - \overline{x})(y_k - \overline{y})}\ ,$$

where $\overline{x}$ and $\overline{y}$ are the mean gene expression of bulk $x$ and pseudobulk $y$, and $x_k$ and $y_k$ are the gene expression value associated with gene $k$ in the bulk or pseudobulk. $\sigma_i = \sigma_{ii}$ and $\sigma_j = \sigma_{jj}$ are the standard deviations within a bulk $i$ or a pseudobulk $j$, respectively. Similar principles apply to cell type proportions of either a sample or a specific cell type across samples.

We used the **Concordance correlation coefficient** (CCC) to measure correlation and agreement of two datasets. It is defined as

$$CCC = \frac{2PCC\sigma_x\sigma_y}{\sigma_x^2\sigma_y^2(\overline{xy})^2}$$

for the agreement of two datasets $X, Y$. The primary application for CCC is to compare samples against a gold-standard. Here, the CCC is used for comparing bulk and pseudobulk expression, ground-truth and predicted proportions for a cell type across samples or a sample across cell types. We found the CCC to penalize predictions that deviate from the ground-truth, while also accounting for a linear relationship between the two variables.

Finally, we computed the **Jensen Shannon Divergence** (JSD), which measures the similarity of two probability distributions $P, Q$. In our case, $P$ is the ground truth bulk sample and $Q$ is the simulated pseudobulk sample. It is based on the Kullback-Leibler divergence $D_{KL}$ that is defined as

$$\text{D}_{\text{KL}} = \sum P * ln\frac{P}{Q}$$

and is asymmetric. To create the JSD as a symmetric variant of $D_{KL}$, first the averaged distribution $M$, defined as

$$\text{M} = \frac{1}{2}(P+Q)$$

is obtained. Then, the $D_{KL}$ of both distributions $P, Q$ against $M$ is calculated and combined using the arithmetic mean. As such, the JSD is defined as

$$\text{JSD} = \frac{1}{2}D_{KL}(P, M) + \frac{1}{2}D_{KL}(Q, M)$$

and thus becomes symmetric. The JSD is in the range $[0, 1]$, whereby two identical distributions have a JSD of 0. We report $1 - JSD$ values in this work.

### 2 Supplementary Figures

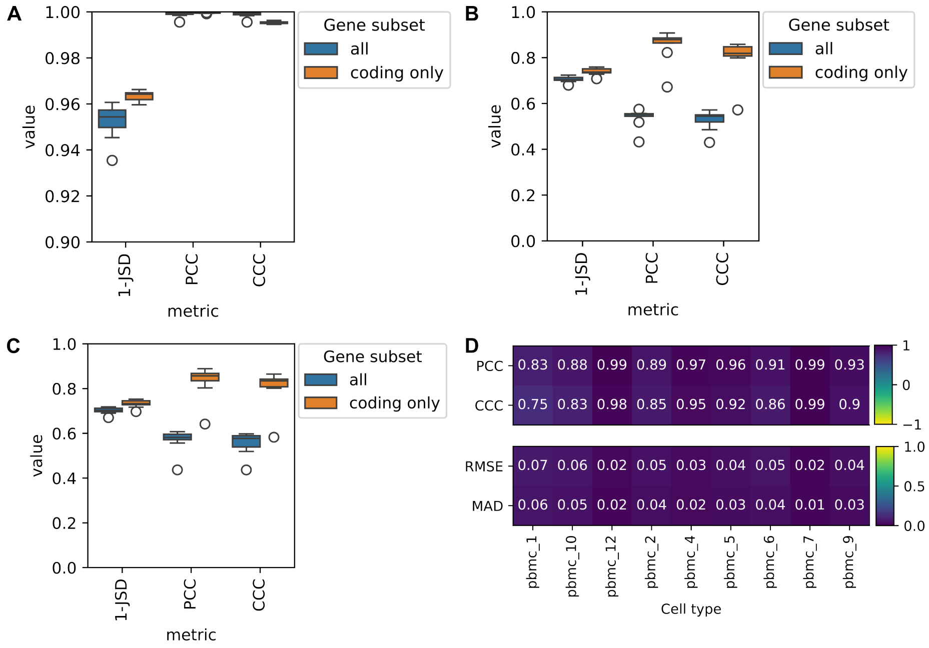


Supplementary Figure S1: **GrooD training with heterogeneous pseudobulks enhances bulk transcriptome deconvolution.**
**A**) Comparison of pseudobulks generated from train (80 %) and test (20 %) data derived from the *Hao* single-cell reference [3] using the same cell type proportions (n=50 pseudobulks compared). Correlation of gene expression was measured on CPM-normalized data for all intersecting genes (all) as well as protein-coding genes (coding only) using 1-JSD, Pearson correlation and CCC as metrics. **B, C**) Comparison of *Finotello* bulk data [2] with *Hao* train data using the underlying cell type proportions from the bulk (B) and without (C) (n=9 samples compared). Again comparisons were performed on CPM-normalized pseudobulk and TPM-normalized bulk data either for all common genes or for coding genes only. **D**) Sample-wise comparison of cell type proportions inferred with GrooD from the *Finotello* bulk data with the ground-truth cell type proportions for n=7 cell types. Both correlation and error metrics are displayed. CCC: Concordance correlation coefficient, PCC: Pearson correlation coefficient, RMSE: root mean squared error, MAD: mean absolute deviation, JSD: Jensen-Shannon divergence.

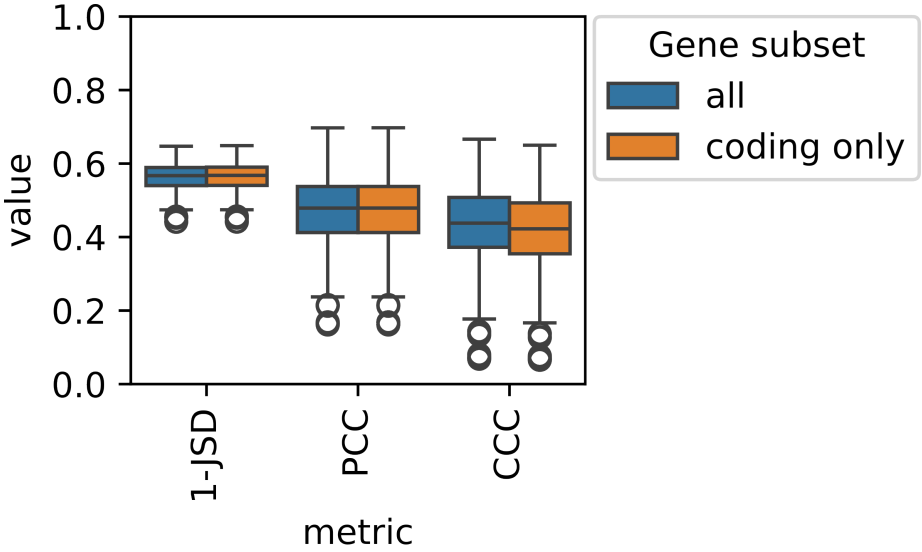


Supplementary Figure S2: **Comparison of *OConnell* bulk data and *Steixner-Kumar* pseudobulks.**
Similarity between *OConnell* bulk RNA-seq data [4] (n=137) and pseudobulks derived from the *Steixner-Kumar* reference [5] (n=137, not considering ground-truth proportions from bulk data) were compared with three quality metrics on all intersecting and only mRNA genes. PCC: Pearson correlation coefficient, CCC: concordance correlation coefficient, JSD: Jensen-Shannon divergence.

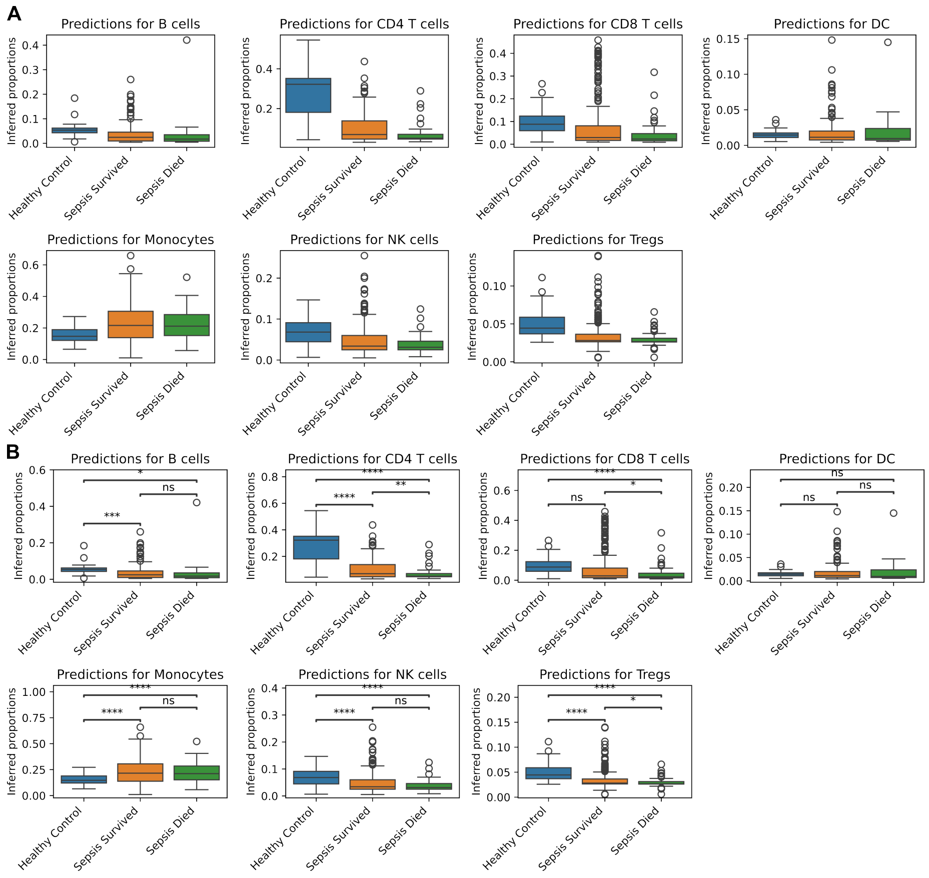


Supplementary Figure S3: **GrooD-inferred cell type proportions across patient subgroups in the *Baghela* sepsis patient cohort.**

**A**) Cell type proportions predicted with GrooD for 389 bulk samples from the Baghela dataset [1] per patient group (healthy control, sepsis survived, sepsis died). This figure complements Figure 3A. **B**) Same plots as in a complemented by statistical tests. Independent t-tests were performed on cell type proportions between patient groups. Asterisks indicate significance (****: $< 1e{-}4$, **: $1e{-}4 < p <= 1e{-}3$, **: $1e{-}3 < p <= 1e{-}2$, *: $1e{-}2 < p <= 1e{-}1$, ns: non-significant). We note that due to the interdependence of cell type proportions independent t-tests, which also do not account for multiple testing correction, are suboptimal means and results are to be treated with caution. We therefore only describe the overall trends in proportions.
